# Plastic but repeatable: rapid adjustments of mitochondrial function and density during reproduction in a wild bird species

**DOI:** 10.1101/708867

**Authors:** Antoine Stier, Pierre Bize, Bin-Yan Hsu, Suvi Ruuskanen

## Abstract

Most of the energy fluxes supporting animal performance flow through mitochondria. Hence, inter-individual differences in performance might be rooted in inter-individual variations in mitochondrial function and density. Furthermore, because the energy required by an individual often changes across life stages, mitochondrial function and density are also expected to show within-individual variation (*i.e.* plasticity). No study so far has repeatedly measured mitochondrial function and density in the same individuals to simultaneously test for within-individual repeatability and plasticity of mitochondrial traits. Here, we repeatedly measured mitochondrial DNA copy number (a proxy of density) and respiration rates from blood cells of female pied flycatchers (*Ficedula hypoleuca*) at the incubation and chick-rearing stages. Mitochondrial density and respiration rates were all repeatable (*R* = [0.45-0.80]), indicating high within-individual consistency in mitochondrial traits across life-history stages. Mitochondrial traits were also plastic, showing a quick (*i.e.* 10 days) down-regulation from incubation to chick-rearing in mitochondrial density, respiratory activity, and cellular regulation by endogenous substrates and/or ATP demand. These downregulations were partially compensated by an increase in mitochondrial efficiency at the chick-rearing stage. Therefore, our study provides clear evidence for both short-term plasticity and high within-individual consistency in mitochondrial function and density during reproduction in a wild bird species.

## Introduction

Mitochondria produce through oxidative phosphorylation more than 90% of the cellular energy fuelling cellular and therefore individual activities [1]. Hence, variation in mitochondrial function and density (*i.e.* amount of mitochondria per cell) have been suggested to account for inter-individual variation in performance [2–4], and in turn individual quality, with high quality individuals consistently outperforming low quality ones. Furthermore, because the energy required by an individual often changes across life stages and contexts (*e.g.* reproduction, hypoxia, cold exposure; [5–7]), mitochondrial function and density are also expected to show within-individual variation (*i.e.* plasticity). Although both hypotheses have received independent support, no study so far has repeatedly measured mitochondrial function and density in the same individuals to simultaneously test for within-individual repeatability and plasticity of mitochondrial traits. Indeed, measures of mitochondrial function almost exclusively rely on invasive sampling that prevents repeated sampling on the same individuals [8].

To fill this knowledge gap, we repeatedly measured mitochondrial function and density from blood cells of free-living female pied flycatchers (*Ficedula hypoleuca*) at the incubation and chick-rearing stages. We previously demonstrated that birds possess functional mitochondria in their red blood cells [9], and thus that mitochondrial function can be repeatedly measured from the same individuals using a minimally invasive repeated blood sampling approach [10]. Hence, our design allowed testing whether mitochondrial traits are repeatable within individuals, with for instance some individuals having consistently more mitochondria with greater respiration rates or efficiencies than others, which is an underlying assumption of the individual quality hypothesis. Furthermore, the reproductive cycle of female birds is well known to be divided into egg-laying, incubation and chick-rearing stages that can differ in their energy constraints [11]. Hence, our design also allowed testing whether mitochondrial traits can quickly respond to changes in energy constraints, supporting the hypothesis that mitochondrial traits are plastic.

## Material & methods

### Fieldwork

Pied flycatcher (*Ficedula hypoleuca*) females breeding in artificial nestboxes in Ruissalo island (Turku, Finland, 60°26.055′N, 22°10.391′E) were captured twice during their reproduction in 2018. Females were captured a first time at day 8 of incubation, and then recaptured at day 7 after hatching (hereafter referred as chick-rearing), leaving a time interval of 10.1 ± 0.1 days between the two sampling occasions. Bird weight was recorded (± 0.1g) and a small blood sample (*i.e.* 25 to 50μL) was taken by puncturating the wing vein with a 26*G* sterile needle and collecting blood using a heparinised capillary. Blood samples were kept cold in the field using ice packs before being transferred to the laboratory for mitochondrial analysis (< 2 hours from blood collection). Blood samples were centrifuged 5min at 2000*g* and plasma was removed before re-suspending the blood cells in 1mL of Mir06 buffer pre-equilibrated at 40°C (see supplementary methods in ESM). Twenty μL of this solution were diluted in 1mL of PBS in order to count the number of cells per sample using an automatic cell counter (Bio-Rad TC20 cell counter).

### Mitochondrial respiration of permeabilized blood cells

We analyzed mitochondrial respiration using a high-resolution respirometry system (Oroboros Instruments, Innsbruck, Austria) adapting the protocol we described in [10] by permeabilizing blood cells in order to get better insights on mitochondrial function (see supplementary methods in ESM and Fig S1). We evaluated 6 mitochondrial respiration rates: 1) *ROUTINE* respiration: endogenous cellular respiration before permeabilization; 2) *complex I* respiration fuelled by exogenous complex I substrates and ADP; 3) *complex I+II* respiration fuelled by exogenous complex I + II substrates and ADP; 4) *complex II* contribution to respiration fuelled by exogenous complex I and II substrates and ADP; 5) *LEAK* respiration due mostly to mitochondrial proton leak (*i.e.* not producing ATP but dissipating heat); and 6) *OXPHOS* respiration that is supporting ATP synthesis through oxidative phosphorylation. We also calculated 3 mitochondrial *flux control ratios* (FCRs), namely FCR_*LEAK/I+II*_ indicating the proportion of mitochondrial respiration being linked to proton leak (*i.e.* an indicator of mitochondrial inefficiency to produce ATP), FCR_*ROUTINE/I+II*_ indicating the proportion of maximal capacity being used under endogenous cellular conditions (*i.e.* reflecting the cellular control of mitochondrial respiration by endogenous ADP/ATP turnover and substrate availability), and FCR_*I/I+II*_ indicating the relative contribution of complex I to total respiration.

### Mitochondrial density

As an indicator of mitochondrial density, we estimated relative mitochondrial DNA copy number (hereafter referred as *mtDNAcn*) by measuring the amount of mitochondrial DNA relative to the nuclear DNA using a qPCR protocol routinely used in humans (*e.g.* [12]) and recently adapted in wild birds [13]. Detailed methodology is available in ESM.

### Statistical analysis

Statistical tests were conducted using *R* 3.4.2. We had measures of body mass and mitochondrial density for 40 females captured both at incubation and chick-rearing. However, due to the strong logistical constraints of working with fresh blood samples for the analysis of mitochondrial respiration, we only had measures of mitochondrial respiration rates for 33 females at incubation and 13 females at the chick-rearing stage. Mitochondrial respiration rates were correlated with mitochondrial density (Pearson correlations: all r > 0.26 and p < 0.10; overall meta-analytic *Zr* = 0.445, p < 0.001, Fig S2). Therefore, we analyzed both cellular respiration rates (O_2_ consumption normalized per cell number, *e.g. ROUTINE*) and respiration rates corrected for mitochondrial density (residuals of the regressions between cellular respirations rates and *mtDNAcn*, *e.g. ROUTINE*_*mt*_) since variations in respiration rates at the cellular level can be explained both by how mitochondria enclosed in those cells are respiring and by the density of mitochondria per cell. Differences between breeding stages were analyzed using paired t-tests and presented as effect size and 95% C.I. following [14]. We evaluated adjusted within-individual repeatability (*i.e.* adjusted for the effects of breeding stage as a fixed factor) and the associated 95% C.I. using the *rptR* package [15].

## Results

We found that body mass, mtDNA copy number, and mitochondrial/cellular respirations rates (*i.e.* after controlling or not for mtDNA copy number), were all significantly decreasing from the incubation to the chick-rearing stage (Fig 1). Females also had more efficient mitochondria during chick rearing (lower FCR_*LEAK/I+II*_) and used less of their mitochondrial maximal capacity (lower FCR_*R/I+II*_), but the relative contribution of complex I to respiration (FCR_*I/I+II*_) was not significantly affected (Fig 1). Despite the major short-term changes observed in mitochondrial traits from incubation to chick-rearing, mitochondrial density and respiration rates were moderately to highly repeatable within an individual, even after accounting for variations in mitochondrial density (Fig 2).

**Fig 1:**
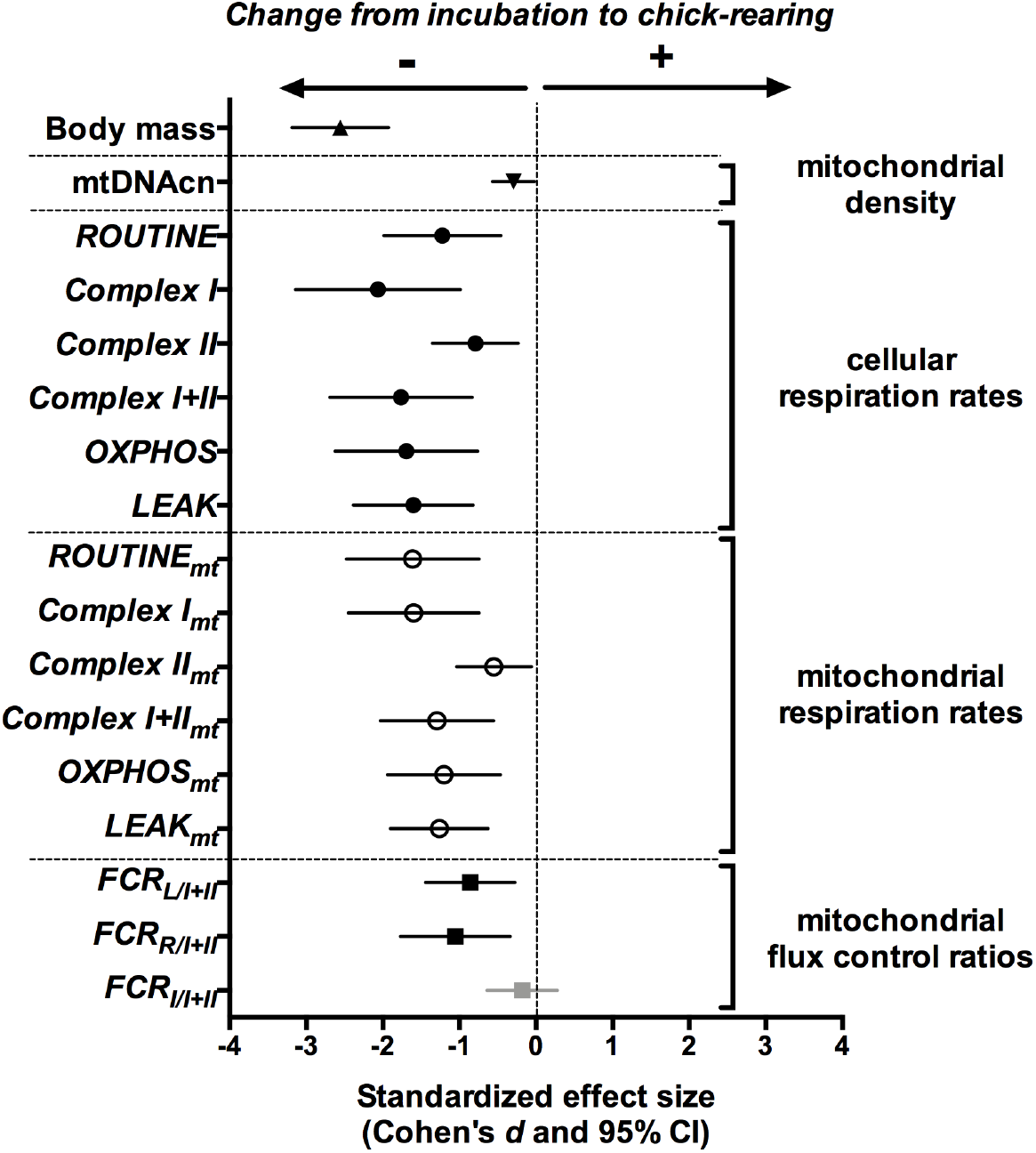
Differences between incubation and chick-rearing stages in body mass, mitochondrial copy number, respiration rates and flux control ratios in female pied flycatchers. Standardized effect size *(Cohen’s d*) are reported with their 95% confidence interval. Significant differences between breeding stages are presented in black and non-significant ones in grey. For respiration rates, we tested both the effects on cellular mitochondrial respiration (*e.g. ROUTINE*), therefore including effects linked both to mitochondrial function and density, and the effects after correcting for mitochondrial density (*i.e.* using regression residuals; labelled *e.g. ROUTINE*_mt_). Detailed information on parameters is given in the method section.

**Fig 2:**
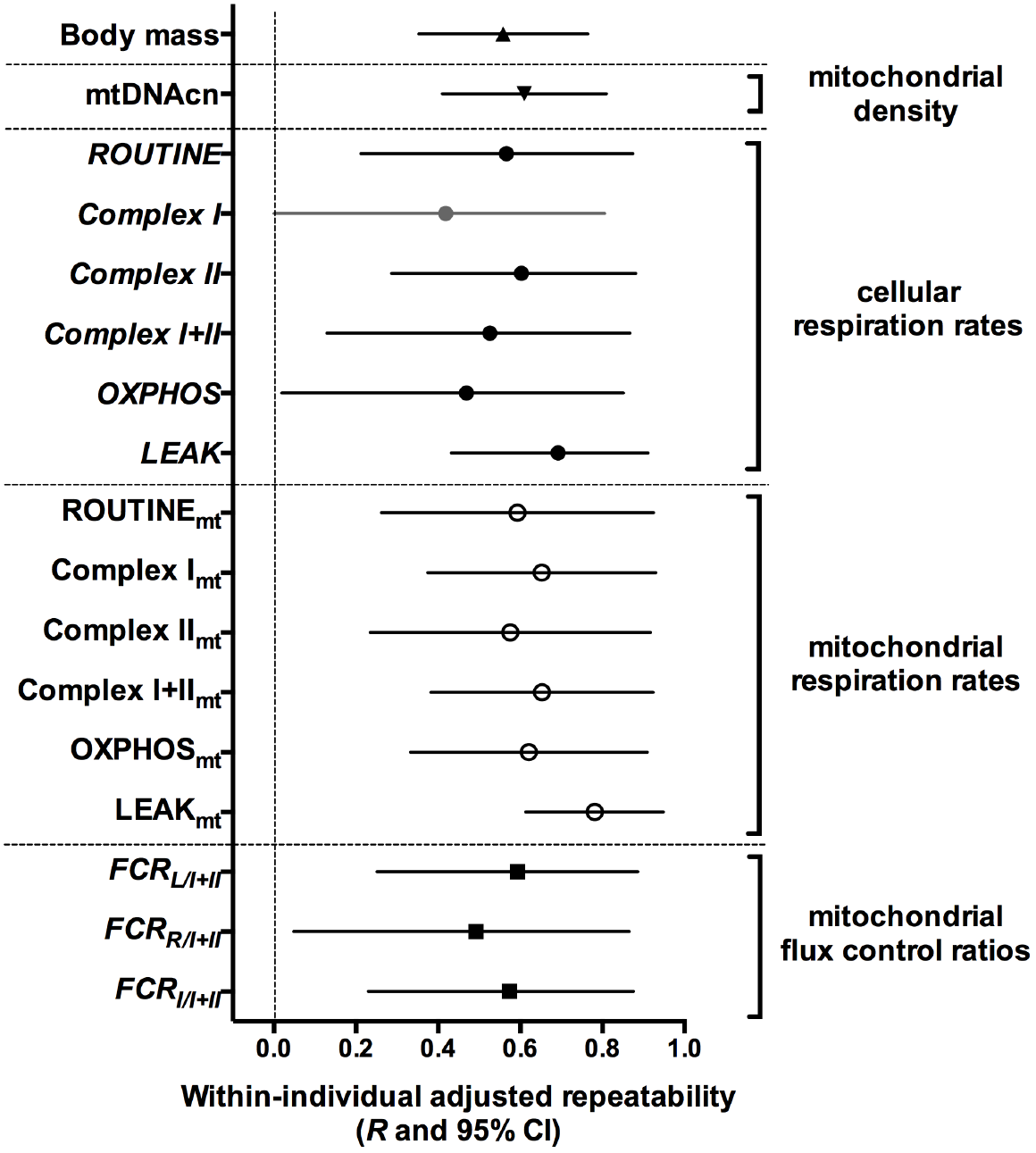
Within-individual adjusted repeatability (*i.e*. consistency) in body mass, mitochondrial density, respiration rates and flux control ratios between incubation and chick-rearing stages in female pied flycatchers. Adjusted repeatability estimates *R* (*i.e.* adjusted for breeding stage fixed effect) are reported with their 95% confidence interval. Significant effects are presented in black and non-significant ones in grey. For mitochondrial respiration rates, we tested both the effects on cellular mitochondrial respiration (*e.g. ROUTINE*), therefore including effects linked both to mitochondrial function and density, and the effects after correcting for mitochondrial density (*i.e.* using regression residuals; labelled *e.g. ROUTINE*_mt_). Detailed information on parameters is given in the method section.

## Discussion

Using repeated measures of mitochondrial traits from the same individual pied flycatchers sampled during incubation and chick-rearing, our study demonstrates that variation in mitochondrial respiration and density is significantly repeatable, as well as significantly plastic within individuals. Individuals with high mitochondrial density and respirations during incubation also had higher values during chick-rearing, and this even if within each individual mitochondrial traits were quickly down-regulated from incubation to chick-rearing. The finding that mitochondrial traits are repeatable provides an important pre-requisite for the hypothesis that repeatable variation in individual performance and quality may be explained by inter-individual variation in mitochondrial traits. Our results also suggest that quick (*i.e.* 10 days) adjustments of cellular bioenergetics can occur during reproduction, most likely in response to differences in energy constraints between breeding stages [11].

Currently, we still know very little about how consistent over time are mitochondrial traits measured in the same individuals, and whether variation in mitochondrial traits measured in one tissue mirrors what is happening in other tissues. We have already shown elsewhere that red blood cell mitochondrial traits are moderately correlated with mitochondrial traits in other tissues ([10] in king penguin, AS unpublished results in Japanese quails), and thus in this study we focused on the first knowledge gap by testing within1 individual repeatability in mitochondrial traits over an interval of 10 days. Our results show that mitochondrial traits were moderately to highly repeatable (mean [min-max] *R* values = 0.63 [0.45-0.80]) within-individuals, to an extent being similar to what we found for female body mass in our study (*R* = 0.58). Interestingly, the repeatability estimates for mitochondrial traits were also in the range of what has been reported by a meta-analysis on whole animal metabolic rates (*R* = 0.57 [16]). Yet, we have to keep in mind that the sampling interval was short (*i.e.* 10 days), and that within-individual repeatability is likely to decrease with increasing duration between sampling points. Significant repeatability could be explained by genetic differences between individuals in genes coding mitochondrial proteins (*e.g.* [17]), but also by potential long-lasting effects of early-life conditions on mitochondrial function [18]. Although within-individual repeatability estimates establish the upper limit for heritability of mitochondrial traits, almost everything remains to be done to understand the relative importance of genetics *vs.* environmental effects in determining mitochondrial traits in wild populations, and to unravel the relationships between mitochondrial and fitness1 related traits. This information is essential to shed light on the importance of mitochondria in shaping variation in individual quality and animal life histories.

Our longitudinal study design also allowed testing for plasticity *per se* in mitochondrial traits by measuring the same individuals under two different environmental conditions rather than, as usually done in the field of mitochondrial biology, by measuring different individuals kept in different environmental conditions. Our results show that both mitochondrial density (*i.e.* estimated as mtDNA copy number) and respiration rates decreased from incubation to chick-rearing, thus suggesting a down-regulation in cellular metabolism, at least in blood cells. The decreases in respiration rates were only moderately explained by changes in mitochondrial density since these decreases remained of moderate to large effect size even after controlling for differences in mitochondrial density. It suggests that female pied flycatchers quickly (10 days) decreased the abundance of respiratory complexes per mitochondria, and to a lesser extent (*i.e.* smaller effect size) the number of mitochondria per cell between incubation and chick-rearing stages. Additionally, mitochondria were also working at a slower pace under endogenous cellular conditions relative to the maximal capacity (*i.e.* lower FCR_*R/I+II*_), suggesting that the control of mitochondrial respiration by endogenous substrates availability and/or ATP demand was also tighter during chick-rearing. Finally, we found a decrease in the relative proton leak (FCR_*L/I+II*_) between incubation and chick-rearing. It suggests that individuals might have increased their mitochondrial efficiency (at least in blood cells), which could potentially carry an oxidative cost [19]. Females are likely more energy-constrained during the chick-rearing stage, and therefore increasing mitochondrial efficiency could be advantageous to maximize short-term performance despite potential delayed costs linked to oxidative stress. The higher relative respiration linked to proton leak during incubation could also potentially be related to the higher need for heat production/dissipation during this breeding stage (*i.e.* to keep the eggs warm) than during chick-rearing. Altogether, our results suggest that mitochondrial adjustments occur at 4 different levels (*i.e.* density, respiration, endogenous regulation and coupling), and thus that studies on mitochondrial function should carefully consider these 4 levels of regulation. Indeed, studies using isolated mitochondria are likely to miss effects related to mitochondrial density and to cellular regulation by endogenous substrates and/or ATP demand. Studies using permeabilized tissues/cells will not be able to tease apart the effects of mitochondrial density and function if not assessing separately mitochondrial density, and could miss effects linked to cellular regulation by endogenous substrates and/or ATP demand if endogenous respiration (*i.e. ROUTINE*) is not assessed. Finally, our study demonstrate that analysing mitochondrial function using blood cells is a promising approach to study the contribution of mitochondrial traits in shaping individual quality and responses to environmental changes.

## Ethics

All procedures were approved by the Animal Experiment Committee of the State Provincial Office of Southern Finland (licence number ESAVI/2902/2018) and by the Environmental Center of Southwestern Finland (license number VARELY549/2018) granted to SR.

## Data availability

Data used in this manuscript is available at: https://doi.org/10.6084/m9.figshare.8960150.v1

## Author’s contribution

AS designed the study, conducted the field and laboratory work, data analysis and wrote the manuscript. SR and BYH conducted the fieldwork, had input on data analysis and commented on the manuscript. PB contributed to the original idea, had input on data analysis and wrote the manuscript.

## Acknowledgements

We are grateful to Lucas Bousseau, Thomas Rossille and Päivi Kotitalo for their help in the field and to the crew of the Marion Dufresnes and the French Polar Institute (IPEV) for hosting AS from 56 to 21° South while writing the manuscript.

## Supplementary methods

### Mitochondrial respiration of permeabilized blood cells

Mitochondrial respiration of permeabilized blood cells was analyzed using a high-resolution respirometry system (Oroboros Instruments, Innsbruck, Austria) and a protocol adapted from Stier et al. (2017, *Methods Ecol Evol*). Blood cells were diluted in respiration buffer Mir06 (0.5 mM EGTA, 3 mM MgCl_2_, 60 mM K-lactobionate, 20 mM taurine, 10 mM KH_2_PO_4_, 20 mM Hepes, 110 mM sucrose, free fatty acid bovine serum albumin (1 g/L), catalase 280U/mL pH 7.1) and added in a closed chamber where O_2_ consumption was recorded following a standard sequential substrate/inhibitor addition protocol. First endogenous respiration of intact blood cells (*i.e. ROUTINE*) was recorded before adding 5 ng.mL^−1^ of digitonin to permeabilize the cells (to allow substrates and ADP to enter the cells). Substrates of complex I (P: pyruvate 5mM and M: malate 2mM) and a saturating amount of ADP (2mM) were added to stimulate mitochondrial respiration fuelled by complex I (hereafter referred as *complex I*). Substrate of complex II (S: succinate 10mM) was then added to stimulate mitochondrial respiration fuelled by both complexes I and II (hereafter referred as *complex I+II*). The difference between these two rates was then calculated to estimate the contribution of complex II to overall respiration (hereafter referred as *complex II*). ATP synthesis was then inhibited with 2.5 μM of oligomycin to estimate mitochondrial inefficiency being mostly linked to proton leak (hereafter referred as *LEAK*). The difference between *complex I+II* and *LEAK* was then calculated to estimate the contribution of oxidative phosphorylation (*i.e.* ATP synthesis) to maximal respiration, hereafter referred as *OXPHOS.* A sequential titration with 0.5μM of the uncoupler FCCP never stimulated respiration above values of *complex I+II* so was removed from the final protocol. Finally, we inhibited mitochondrial respiration using antimycin A (2.5μM), and this residual non-mitochondrial O_2_ consumption was subtracted from the mitochondrial parameters described above. Respiration rates were expressed as pmol O_2_.s^−1^.10^6^ cells^−1^. Oxygen levels within the chamber were maintained between 140 and 200μM of O_2_ using 0.5μL injections of 200mM H_2_O_2_ between titration steps. Mitochondrial responses to this chemical titration is presented in ESM Fig S1. We also calculated 3 mitochondrial *flux control ratios* (FCRs), namely FCR_*L/I+II*_ indicating the proportion of mitochondrial respiration being linked to proton leak (*i.e.* an indicator of mitochondrial inefficiency to produce ATP), FCR_*R/I+II*_ indicating the proportion of maximal capacity being used under endogenous cellular conditions (*i.e.* reflecting the cellular control of mitochondrial respiration by endogenous ADP/ATP turnover and substrate availability), and FCR_*I/I+II*_ indicating the relative contribution of complex I to total respiration. The low amount of blood being available with such small birds prevented us to estimate the technical repeatability in this study, but our previous results in king penguins revealed a high repeatability of the method (Stier et al. 2017, *Methods Ecol Evol*), and the high within-individual repeatability estimates reported here in the main text (Fig 2) confirms that our technical repeatability was high although not directly evaluated.

### Mitochondrial density

As an indicator of mitochondrial density, we estimated relative mtDNA copy number by measuring the amount of mitochondrial DNA relative to the nuclear DNA (hereafter referred as *mtDNAcn*) by qPCR on a 384-QuantStudio™ 12K Flex Real-Time PCR System (Thermo Fisher). DNA was extracted (using Macherey-Nagel Blood QuickPure spin columns) from blood samples and diluted to 1.2ng.μL^−1^. We used RAG1 as a nuclear single-copy control gene (verified using a BLAST analysis on the collared flycatcher *Ficedula albicollis* genome; F: GCAGATGAACTGGAGGCTATAA, R: CAGCTGAGAAACGTGTTGATTC). We used cytochrome oxidase subunit 2 (COI2) as a specific mitochondrial gene after verifying that it was not duplicated as a pseudo-gene in the nuclear genome using a BLAST analysis on the collared flycatcher genome (F: GGAGACGACCAAGTCTACAATG; R: TTTCCGAACCCTCCGATTATG). Melt curve and electrophoresis analyses confirmed that a single amplicon of the expected length was generated by PCR with the designed primers. For the qPCR assay, the reactions were performed in a total volume of 12μL including 6ng of DNA, primers at a final concentration of 200nM and 6μL of Absolute Blue qPCR Mix SYBR Green low ROX (Thermo Scientific). RAG1 and COI2 reactions were performed in triplicates on the same plates (4 plates in total); the qPCR conditions were: 15 min at 95°C, followed by 40 cycles of 15 s at 95°C, 30 s at 60°C and 30s at 72°C. A DNA sample being a pool of DNA from 10 individuals was used as a reference sample and was included in triplicate on every plate. The efficiency of each amplicon was estimated from a standard curve of the reference sample ranging from 1.5 to 24ng. The mean reaction efficiencies were 106.8 ± 1.4% for RAG1 and 91.1 ± 0.7% for COI2. The relative mtDNA copy number (*mtDNAcn*) of each sample was calculated as (1+*Ef*_COI2_)^ΔCqCOI2^/(1+*Ef*_RAG1_)^ΔqRAG1^; *Ef* being the amplicon efficiency, and ΔCq the difference in Cq-values between the reference sample and the focal sample. Intra-plate technical repeatability of *mtDNAcn* based on triplicates was 0.81 (95% C.I. [0.76-0.85]), and the inter-plate technical repeatability based on two repeated plates was 0.92 (95% C.I. [0.88-0.95]).

## Supplementary figures

**Fig S1:**
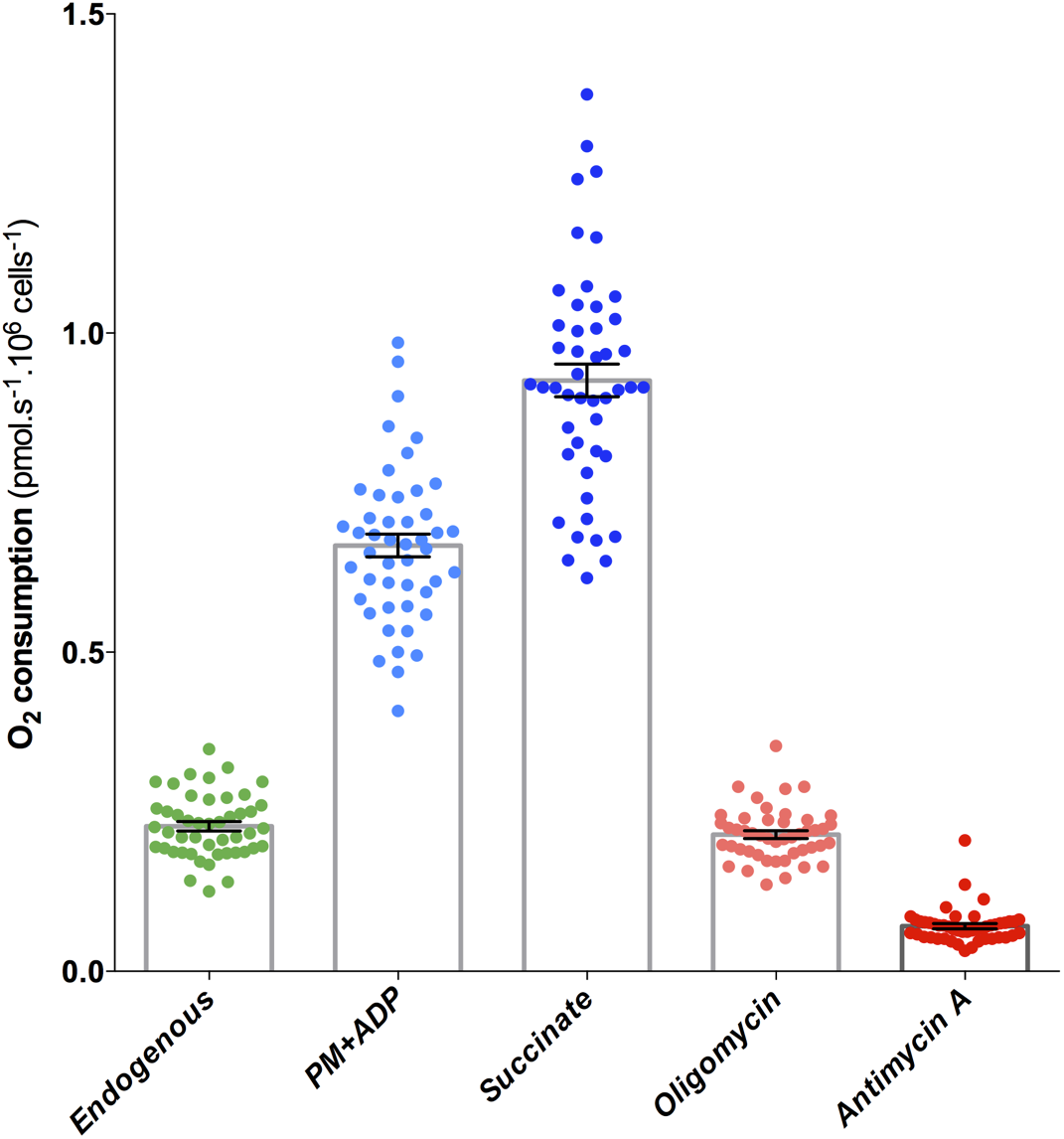
*In-vitro* cellular O_2_ consumption of pied flycatcher red blood cells in response to a standard sequential substrate/inhibitor addition protocol. Cells have been permeabilized with 5ng.mL^−1^ of digitonin after recording the endogenous respiration of intact cells. Details about chemical additions are given in the supplementary methods described above. Data are presented as individual data points and mean ± SE.

**Fig S2:**
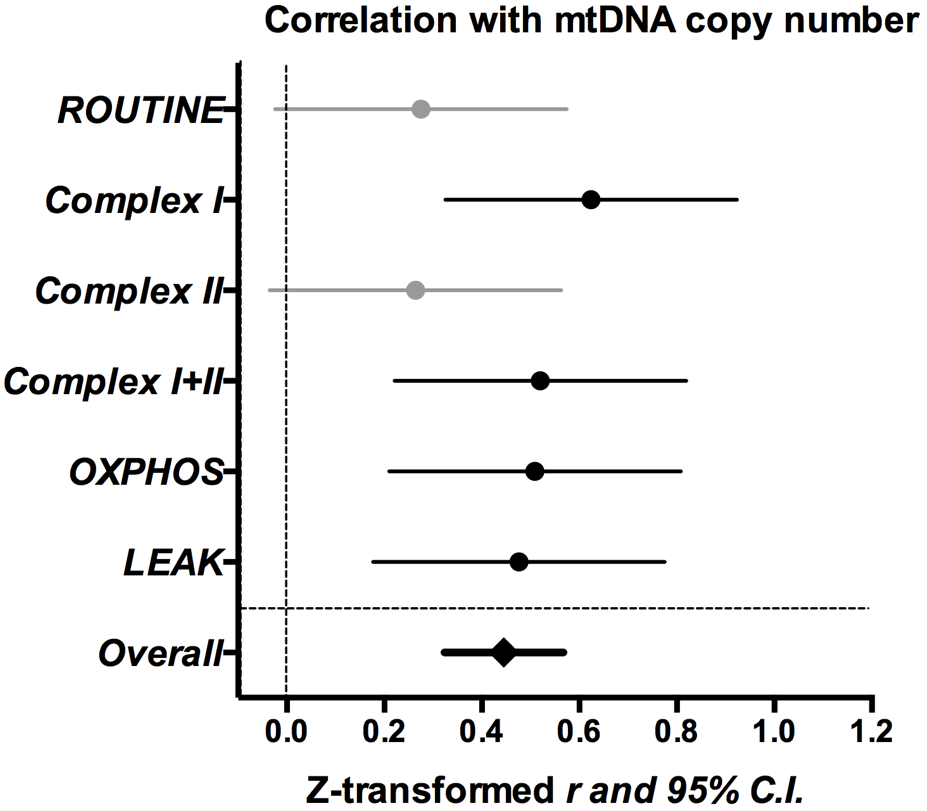
Pearson’s correlations between cellular respiration rates and mitochondrial DNA copy number (*mtDNAcn*) measured in blood cells of female pied flycatchers. A metaJanalytic overall correlation has been calculated using the OpenMEE software. Z-transformed *r* are presented (instead of *r* due to meta-analysis statistical constraints), along with their 95% confidence intervals. Details about mitochondrial parameters are presented in the supplementary methods above. Significant parameters are shown in black and non-significant ones (but p < 0.10) in grey. Data were collected on 33 females at incubation and 13 females at chick rearing.

## References

1. Nicholls, D. G. & Ferguson, S. J. 2002 Bioenergetics. Academic press

2. Stier, A., Bize, P., Roussel, D., Schull, Q., Massemin, S. & Criscuolo, F. 2014 Mitochondrial uncoupling as a regulator of life-history trajectories in birds: an experimental study in the zebra finch. The Journal of Experimental Biology 217, 3579–3589. (doi:10.1242/jeb.103945)

3. Salin, K., Auer, S. K., Rey, B., Selman, C. & Metcalfe, N. In press. Variation in the link between oxygen consumption and ATP production, and its relevance for animal performance. Proceedings of the Royal Society B: Biological Sciences 282. (doi:http://doi.org/10.1098/rspb.2015.1028)

4. Hood, W. R. et al. 2018 The Mitochondrial Contribution to Animal Performance, Adaptation, and Life-History Variation. Integrative and Comparative Biology 58, 480–485. (doi:10.1093/icb/icy089)

5. Toyomizu, M., Ueda, M., Sato, S., Seki, Y., Sato, K. & Akiba, Y. 2002 Cold-induced mitochondrial uncoupling and expression of chicken UCP and ANT mRNA in chicken skeletal muscle. FEBS Letters 529, 313–318.

6. Sussarellu, R., Dudognon, T., Fabioux, C., Soudant, P., Moraga, D. & Kraffe, E. 2013 Rapid mitochondrial adjustments in response to short-term hypoxia and re1 oxygenation in the Pacific oyster, Crassostrea gigas. The Journal of Experimental Biology 216, 1561–1569. (doi:10.1242/jeb.075879)

7. Hyatt, H. W., Zhang, Y., Hood, W. R. & Kavazis, A. N. 2017 Changes in Metabolism, Mitochondrial Function, and Oxidative Stress Between Female Rats Under Nonreproductive and 3 Reproductive Conditions. Reproductive Sciences 26, 114–127. (doi:10.1177/1933719118766264)

8. Stier, A., Reichert, S., Criscuolo, F. & Bize, P. 2015 Red blood cells open promising avenues for longitudinal studies of ageing in laboratory, non-model and wild animals. Experimental Gerontology 71, 118–134. (doi:10.1016/j.exger.2015.09.001)

9. Stier, A. et al. 2013 Avian erythrocytes have functional mitochondria, opening novel perspectives for birds as animal models in the study of ageing. Frontiers in Zoology. 10, 33. (doi:10.1186/174219994110133)

10. Stier, A., Romestaing, C., Schull, Q., Lefol, E., Robin, J.-P, Roussel, D. & Bize, P. 2017 How to measure mitochondrial function in birds using red blood cells: a case study in the king penguin and perspectives in ecology and evolution. Methods in Ecology and Evolution 8, 1172–1182. (doi:10.1111/20411210X.12724)

11. Deeming, D. C. & Reynolds, S. J. 2015 Nests, eggs, and incubation: new ideas about avian reproduction. Oxford Scholarship Online. (DOI:10.1093/acprof:oso/9780198718666.001.0001)

12. Mengel-From, J., Thinggaard, M., Dalgård, C., Kyvik, K. O., Christensen, K. & Christiansen, L. 2014 Mitochondrial DNA copy number in peripheral blood cells declines with age and is associated with general health among elderly. Hum. Genet. 133, 1149–1159. (doi:10.1007/s0043910141145819)

13. Velando, A., Noguera, J. C., da Silva, A. & Kim, S.-Y 2019 Redox-regulation and life1 history trade-offs: scavenging mitochondrial ROS improves growth in a wild bird. Scientific Reports. 9, 1–9. (doi:10.1038/s41598101913853515)

14. Nakagawa, S. & Cuthill, I. C. 2007 Effect size, confidence interval and statistical significance: a practical guide for biologists. Biological Reviews 82, 591–605. (doi:10.1111/j.14691185X.2007.00027.x)

15. Stoffel, M. A., Nakagawa, S. & Schielzeth, H. 2017 rptR: repeatability estimation and variance decomposition by generalized linear mixed-effects models. Methods in Ecology and Evolution 8, 1639–1644. (doi:10.1111/20411210X.12797)

16. Nespolo, R. F. & Franco, M. 2007 Whole-animal metabolic rate is a repeatable trait: a meta-analysis. The Journal of Experimental Biology 210, 3877–3878. (doi:10.1242/jeb.013110)

17. Toews, D. P. L., Mandic, M., Richards, J. G. & Irwin, D. E. 2013 Migration, mitochondria, and the yellow-rumped warbler. Evolution 68, 241–255. (doi:10.1111/evo.12260)

18. Dillin, A. 2002 Rates of Behavior and Aging Specified by Mitochondrial Function During Development. Science 298, 2398–2401. (doi:10.1126/science.1077780)

19. Salin, K., Villasevil, E. M., Anderson, G. J., Auer, S. K., Selman, C., Hartley, R. C., Mullen, W., Chinopoulos, C. & Metcalfe, N. B. 2018 Decreased mitochondrial metabolic requirements in fasting animals carry an oxidative cost. Functional Ecology 32, 2149–2157. (doi:10.1111/136512435.13125)

